# Robust deep learning based protein sequence design using ProteinMPNN

**DOI:** 10.1101/2022.06.03.494563

**Authors:** J. Dauparas, I. Anishchenko, N. Bennett, H. Bai, R. J. Ragotte, L. F. Milles, B. I. M. Wicky, A. Courbet, R. J. de Haas, N. Bethel, P. J. Y. Leung, T. F. Huddy, S. Pellock, D. Tischer, F. Chan, B. Koepnick, H. Nguyen, A. Kang, B. Sankaran, A. K. Bera, N. P. King, D. Baker

## Abstract

While deep learning has revolutionized protein structure prediction, almost all experimentally characterized de novo protein designs have been generated using physically based approaches such as Rosetta. Here we describe a deep learning based protein sequence design method, ProteinMPNN, with outstanding performance in both in silico and experimental tests. The amino acid sequence at different positions can be coupled between single or multiple chains, enabling application to a wide range of current protein design challenges. On native protein backbones, ProteinMPNN has a sequence recovery of 52.4%, compared to 32.9% for Rosetta. Incorporation of noise during training improves sequence recovery on protein structure models, and produces sequences which more robustly encode their structures as assessed using structure prediction algorithms. We demonstrate the broad utility and high accuracy of ProteinMPNN using X-ray crystallography, cryoEM and functional studies by rescuing previously failed designs, made using Rosetta or AlphaFold, of protein monomers, cyclic homo-oligomers, tetrahedral nanoparticles, and target binding proteins.

**One-sentence summary:** A deep learning based protein sequence design method is described that is widely applicable to current design challenges and shows outstanding performance in both in silico and experimental tests.

## Main text

The protein sequence design problem is to find, given a protein backbone structure of interest, an amino acid sequence that will fold to this structure. Physically based approaches like Rosetta approach sequence design as an energy optimization problem, searching for the combination of amino acid identities and conformations that have the lowest energy for a given input structure. Recently deep learning approaches have shown considerable promise in rapidly generating plausible amino acid sequences given monomeric protein backbones without need for compute intensive explicit consideration of sidechain rotameric states (*1-6*). However, the methods described thus far are limited in their applicability to the wide range of current protein design challenges, and have not been extensively validated experimentally.

We set out to develop a deep learning based protein sequence design method broadly applicable to design of monomers, cyclic oligomers, protein nanoparticles, and protein-protein interfaces. We began from a previously described message passing neural network (MPNN) with 3 encoder and 3 decoder layers and 128 hidden dimensions which predicts protein sequences in an autoregressive manner from N to C terminus using protein backbone features – distances between Ca-Ca atoms, relative Ca-Ca-Ca frame orientations and rotations, and backbone dihedral angles–as input (*1*). We first sought to improve performance of the model on recovering the amino acid sequences of native monomeric proteins given their backbone structures, using as training and validation sets 19.7k high resolution single-chain structures from the PDB split based on the CATH (*7*) protein classification (see Methods). We experimented with adding distances between N, Ca, C, O and a virtual Cb placed based on the other backbone atoms as additional input features, hypothesising that they would enable better inference than backbone dihedral angle features. This resulted in a sequence recovery increase from 41.2% (baseline model) to 49.0% (experiment 1), see Table 1 below; interatomic distances evidently provide a better inductive bias to capture interactions between residues than dihedral angles or N-Ca-C frame orientations. We next experimented with introducing edge updates in addition to the node updates in the backbone encoder neural network (experiment 2). Combining additional input features and edge updates leads to a sequence recovery of 50.5% (experiment 3). To determine the range over which backbone geometry influences amino acid identity, we experimented with 16, 24, 32, 48, and 64 nearest Ca neighbor neural networks (Figure S1A), and found that performance saturated at 32-48 neighbors. Unlike the protein structure prediction problem, locally connected graph neural networks can be used to model the structure to sequence mapping problem because protein backbones provide a notion of local neighborhoods which primarily determine sequence identities.

**Table 1.** Single chain sequence design performance on CATH held out test split. Test accuracy (percentage of correct amino amino acids recovered) and test perplexity (exponentiated categorical cross entropy loss per residue) are reported for models trained on the native backbone coordinates (left, normal font) and models trained with Gaussian noise (std=0.02Å) added to the backbone coordinates (right, bold font); all test evaluations are with no added noise. The final column shows sequence recovery on 5,000 AlphaFold protein backbone models with average pLDDT > 80.0 randomly chosen from UniRef50 sequences.

| Noise level when training: 0.00Å/**0.02Å** | Modification | Number of Parameters | PDB Test Accuracy | PDB Test Perplexity | AlphaFold Model Accuracy |
| --- | --- | --- | --- | --- | --- |
| Baseline model | None | 1.381 mln | 41.2/**40.1** | 6.51/**6.77** | 41.4/**41.4** |
| Experiment 1 | Add N, Ca, C, Cb, O distances | 1.430 mln | 49.0/**46.1** | 5.03/**5.54** | 45.7/**47.4** |
| Experiment 2 | Update encoder edges | 1.629 mln | 43.1/**42.0** | 6.12/**6.37** | 43.3/**43.0** |
| Experiment 3 | Combine 1 and 2 | 1.678 mln | 50.5/**47.3** | 4.82/**5.36** | 46.3/**47.9** |
| Experiment 4 | Experiment 3 with random instead of forward decoding | 1.678 mln | 50.8/**47.9** | 4.74/**5.25** | 46.9/**48.5** |

To enable application to a broad range of single and multi-chain design problems, we replaced the fixed N to C terminal decoding order with an order agnostic autoregressive model in which the decoding order is randomly sampled from the set of all possible permutations (*8*). This also resulted in a modest improvement in sequence recovery (Table 1, experiment 4). Order agnostic decoding enables design in cases where, for example, the middle of the protein sequence is fixed and the rest needs to be designed, as in protein binder design where the target sequence is known; decoding skips the fixed regions but includes them in the sequence context for the remaining positions (Figure 1B). For multi-chain design problems, to make the model equivariant to the order of the protein chains, we kept the per chain relative positional encoding capped at ±32 residues (*9*) and added a binary feature indicating if the interacting pair of residues are coming from the same or different chains.

**Fig. 1.**
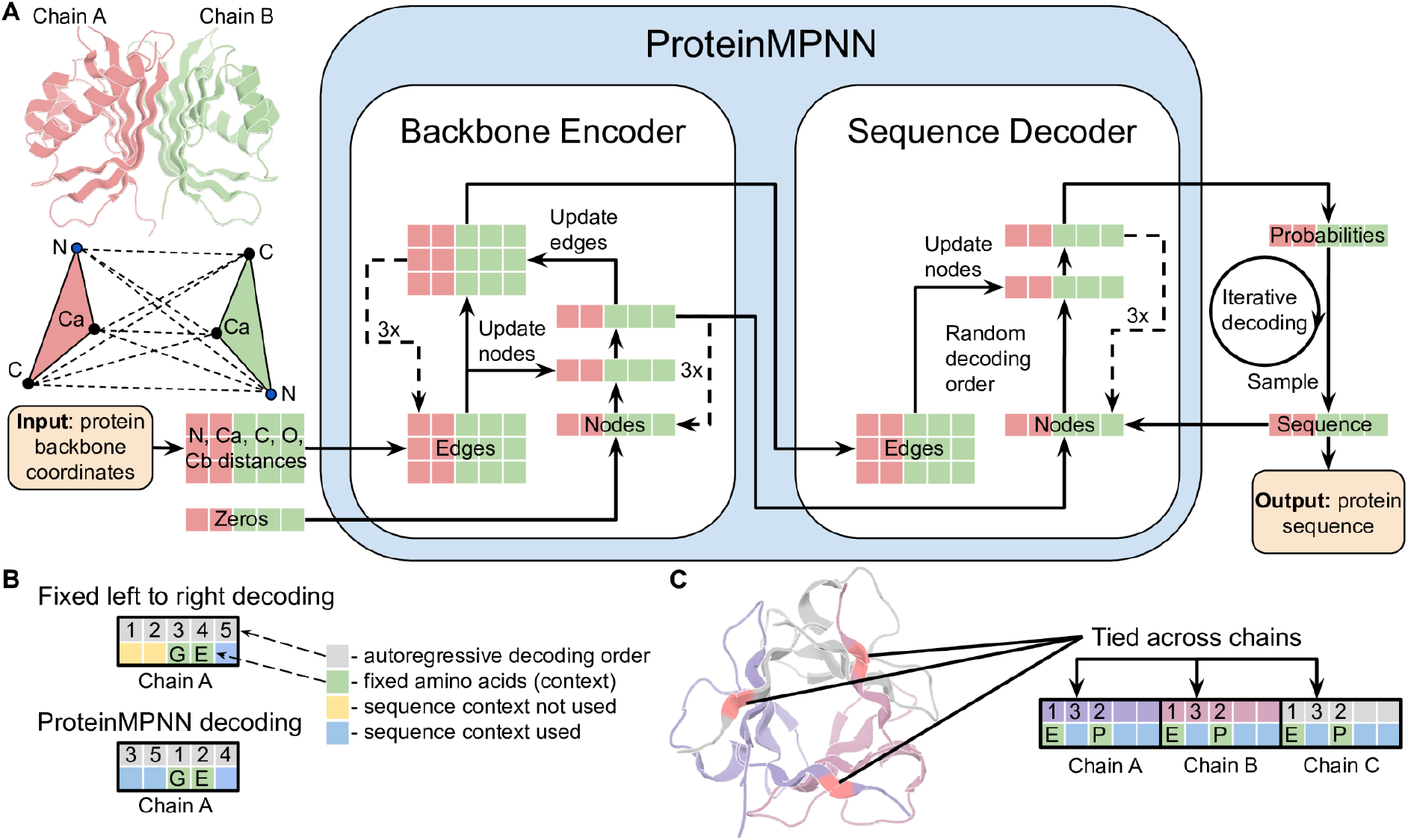
ProteinMPNN architecture. (**A**) Distances between N, Ca, C, O, and virtual Cb are encoded and processed using a message passing neural network (Encoder) to obtain graph node and edge features. The encoded features together with a partial sequence are used to generate amino acids iteratively in a random decoding order. (**B**) A fixed left to right decoding cannot use sequence context (green) for preceding positions (yellow) whereas a model trained with random decoding orders can be used with arbitrary decoding order during the inference. The decoding order can be chosen such that the fixed context is decoded first. (**C**) Residue positions within and between chains can be tied together, enabling symmetric, repeat protein, and multistate design. In this example, a homo-trimer is designed with coupling of positions in different chains. Predicted logits for tied positions are averaged to get a single probability distribution from which amino acids are sampled.

We took advantage of the flexibility of the decoding order, which enables selection during inference of a decoding order appropriate for the specific task, to enable the fixing of residue identities in sets of corresponding positions (the residues at these positions are decoded at the same time). For example, for a C2 homodimer backbone with two chains A and B with sequence A1, A2, A3,.. and B1, B2, B3,…, the amino acids for chains A and B have to be the same for corresponding indices; we implement this by predicting logits (unnormalized probabilities) for A1 and B1 first and then combine these two predictions to construct a normalized probability distribution from which a joint amino acid is sampled (Figure 1C). For pseudosymmetric sequence design, residues within, or between chains can be similarly constrained; for example for repeat protein design, the sequence in each repeat unit can be kept fixed. Multi-state protein sequence design to generate a single sequence that encodes two or more desired states can be achieved by predicting logits for each state and then averaging; more generally a linear combination of predicted logits with some positive and negative coefficients can be used to upweight, or downweight specific backbone states to achieve explicit positive-negative sequence design. The architecture of this multichain and symmetry aware (positionally coupled) model, which we call ProteinMPNN, is outlined schematically in Figure 1A. We trained ProteinMPNN on protein assemblies in the PDB (as of Aug 02, 2021) determined by X-ray crystallography or cryoEM to better than 3.5Å resolution and with less than 10,000 residues. Sequences were clustered at 30% sequence identity cutoff using mmseqs2 (*10*) resulting in 25,361 clusters (see Methods).

For a test set of 402 monomer backbones we redesigned sequences using Rosetta fixed backbone combinatorial sequence design (one round of the PackRotamersMover (*11-12*) with default options and the beta_nov16 score function) and ProteinMPNN. Although requiring only a small fraction of the compute time (1.2 seconds versus 4.3 minutes for 100 residues), ProteinMPNN had a much higher overall native sequence recovery (52.4% vs 32.9%), with improvements across the full range of residue burial from protein core to surface (Figure 2A). Differences between designed and native amino acid biases for the core, boundary and surface regions for the two methods are shown in Figure S2.

**Fig. 2.**
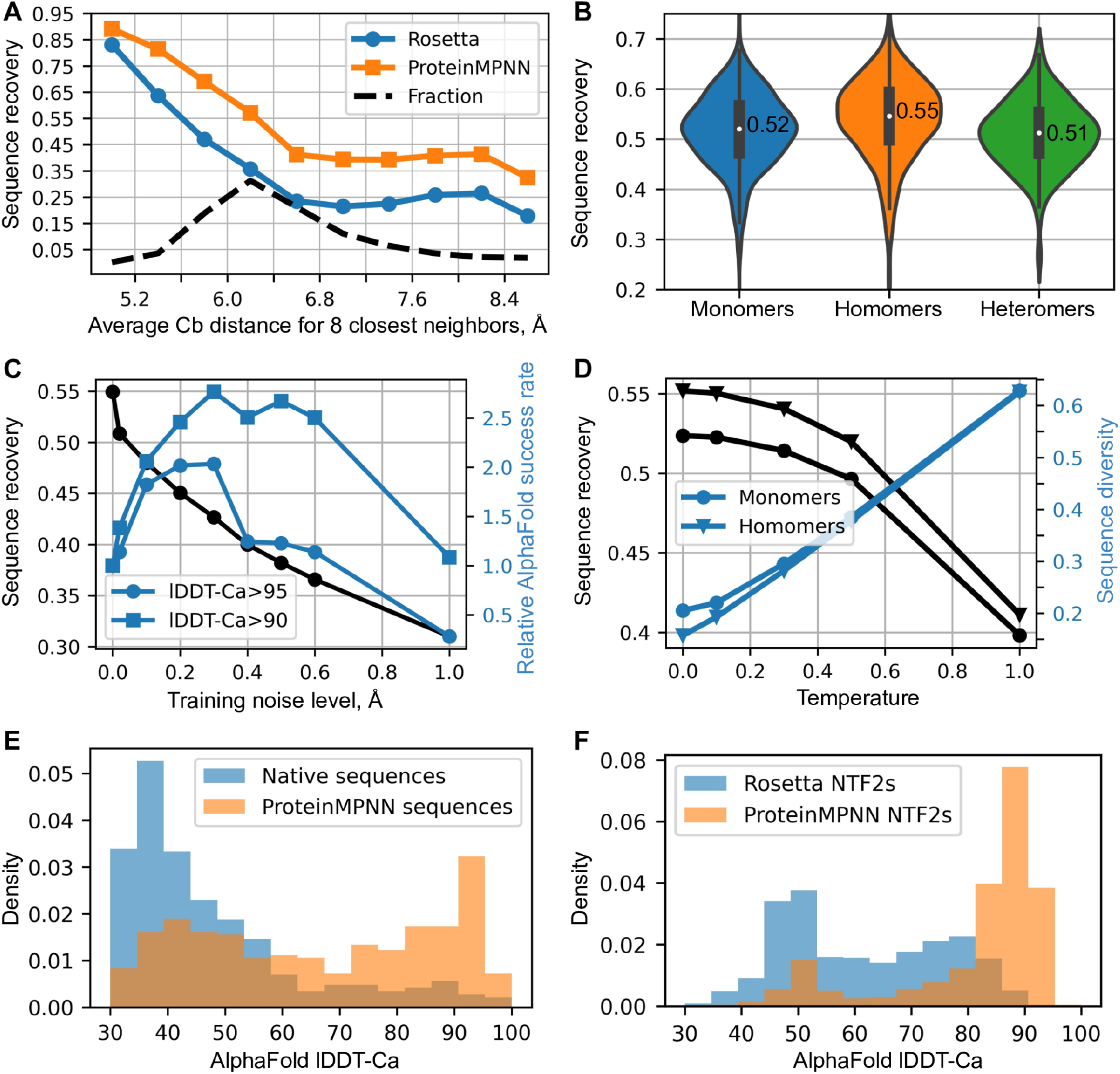
In silico evaluation of ProteinMPNN. (**A**) ProteinMPNN has higher native sequence recovery than Rosetta. The average Cb distance of the 8 closest neighbors (x axis) reports on burial, with most buried positions on the left and more exposed on the right; ProteinMPNN outperforms Rosetta at all levels of burial. Average sequence recovery for ProteinMPNN was 52.4%, compared to 32.9% for Rosetta. (**B**) ProteinMPNN has similarly high sequence recovery for monomers, homo-oligomers, and hetero-oligomers; violin plots are for 690 monomers, 732 homomers, 98 heteromers. (**C**) Sequence recovery (black) and relative AlphaFold success rates (blue) as a function of training noise level. For higher accuracy predictions (circles) smaller amounts of noise are optimal (1.0 corresponds to 1.8% success rate), while to maximize prediction success at a lower accuracy cutoff (squares), models trained with more noise are better (1.0 corresponds to 6.7% success rate). (**D**) Sequence recovery and diversity as a function of sampling temperature. Redesign of native protein backbones with ProteinMPNN considerably increases AphaFold prediction accuracy compared to the original native sequence using no multiple sequence information. Single sequences (designed or native) were input in both cases. (**F**) ProteinMPNN redesign of previous Rosetta designed NTF2 fold proteins (3,000 backbones in total) results in considerably improved AlphaFold single sequence prediction accuracy.

We evaluated ProteinMPNN on a test set of 690 monomers, 732 homomers (with less than 2000 residues), and 98 heteromers. The median overall sequences recoveries were 52% for monomers, 55% for homomers, and 51% for heteromers (Figure 2B). In all three cases, sequence recovery correlated closely with residue burial ranging from 90-95% in the deep core to 35% on the surface (Figure S1B): the amount of local geometric context determines how well residues can be recovered at specific positions. For homomers, we found best results with averaging logits between symmetry related positions: unconstrained design without symmetry, averaged probabilities, and averaged logits resulted in 52%, 53%, and 55% median sequence recoveries respectively (Figure S1C). Because of the non-local context, sequence recovery is no longer a monotonic function of the average Cb neighbor distance; some residues get information from their symmetric counterparts via averaging of probabilities (Figure S1B).

### Training with backbone noise improves model performance for protein design

While recent protein sequence design approaches have focused on maximizing native sequence recovery, this is not necessarily optimal for actual protein design applications. Native sequence recovery is likely highest for models trained on perfect protein backbones, and with stochastic sequence inference carried out at low temperature. We reasoned, however, that improved protein design performance might be achieved by models trained with backbone noise and with inference conducted at higher temperature, as described in the following paragraphs.

Robustness to small displacements in atomic coordinates is a desirable feature for sequence design methods in real world applications where the protein backbone geometry is not known at atomic resolution. We found that training models on backbones to which Gaussian noise (std=0.02Å) had been added improved sequence recovery on confident protein structure models generated by AlphaFold (average pLDDT>80.0) from UniRef50, while the sequence recovery on unperturbed PDB structures decreased as expected (Table 1). Crystallographic refinement may impart some memory of amino acid identity in the backbone coordinates which is recovered by the model trained on perfect backbones but not present in predicted structures; since the goal is to identify optimal sequences given the overall backbone context the more robust model is preferable.

AlphaFold (*9*) and RoseTTAfold (*13*) produce remarkably good structure predictions for native proteins given multiple sequence alignments which can contain substantial co-evolutionary and other information reflecting aspects of the 3D structure, but generally produce much poorer structures when provided only with a single sequence. We reasoned that ProteinMPNN might generate sequences for native backbones more strongly encoding the structures than the original native sequences, as evolution in most cases does not optimize for stability, and completely redesigned a set of 396 native structures. We found in single sequence AlphaFold predictions that ProteinMPNN sequences were predicted to adopt the original native backbone structures much more confidently and accurately than the original native sequences (Figure 2E). We also tested ProteinMPNN on a set of de novo designed scaffolds which contain a wide range of ligand binding pockets. Whereas only a small fraction of the original Rosetta designed sequences were predicted to fold to the design target structures, following ProteinMPNN redesign the majority were confidently predicted to fold to close to the design target structures (Figure 2F). This should substantially increase the utility of these scaffolds for design of protein binding and enzymatic functions–the likelihood that the sequences fold to the desired structures is higher, and designed enzymes and small molecule binding proteins based on these scaffolds can be evaluated using similar structure prediction tests prior to experimental characterization.

We found that the strength of the single sequence to structure mapping, as assessed by AlphaFold, was higher for models trained with additional backbone noise. As noted above, the average sequence recovery for perfect backbones decreases with increasing amounts of noise added during training (Figure 2C) as these models are not able to pick up on fine details of the backbone geometry. In contrast, sequences encoded by noised ProteinMPNN models are more accurately decoded into 3D coordinates by AlphaFold, likely because noised models focus more on overall topological features than fine local structural details (which are blurred during noising). For example, a model trained with 0.3Å noise generated 2-3 times more sequences with AlphaFold predictions within lDDT-Ca (*14*) of 95.0 and 90.0 of the true structures than unnoised or slightly noised models (Figure 2C). In protein design calculations, the models trained with larger amounts of noise have the advantage of generating sequences which more strongly map to the target structures by prediction methods (this increases frequency of designs passing prediction based filters, and may correspondingly also increase the frequency of actual folding to the desired target structure).

Because the sequence determinants of protein expression, solubility and function are not perfectly understood, in most protein design applications it is desirable to test multiple designed sequences experimentally. We found that the diversity of sequences generated by MPNN could be considerably increased, with only a very small decrease in average sequence recovery, by carrying out inference at higher temperatures (Figure 2D). We also found that a measure of sequence quality derived from the ProteinMPNN, the averaged log probability of the sequence given the structure, correlated strongly with native sequence recovery over a range of temperatures (Figure S3A), enabling rapid ranking of sequences for selection for experimental characterization.

### Experimental evaluation of ProteinMPNN

While in silico native protein sequence recovery is a useful benchmark, the ultimate test of a protein design method is its ability to generate sequences which fold to the desired structure and have the desired function when tested experimentally. We evaluated ProteinMPNN on a representative set of design challenges ranging from protein monomer design, protein nanocage design, and protein function design. In each case, we attempted to rescue previous failed designs with sequences generated using Rosetta or AlphaFold–we kept the backbones of the original designs fixed but discarded the original sequences and generated new ones using ProteinMPNN. Synthetic genes encoding the designs were obtained, and the proteins expressed in *E. coli* and characterized biochemically and structurally.

We first tested the ability of ProteinMPNN to design amino acid sequences for protein backbones generated by deep network hallucination using AlphaFold (AF). Starting from a random sequence, a Monte Carlo trajectory is carried out optimizing the extent to which AF predicts the sequence to fold to a well-defined structure (see accompanying paper for details, *Wicky et al*.). These calculations generated a very wide range of protein sequences and backbones for both monomers and oligomers that differ considerably from those of native structures. In initial tests, the sequences generated by AF were encoded in synthetic genes, and we attempted to express 150 proteins in *E. coli*. However, we found that the AF generated sequences were mostly insoluble (median soluble yield: 9 mg per liter of culture equivalent Figure 3A). To determine whether ProteinMPNN could overcome this problem, we generated sequences for a subset of these backbones with ProteinMPNN; residue identities at symmetry-equivalent positions were tied by averaging logits as described above. The designed sequences were again encoded in synthetic genes and the proteins produced in *E. coli*. The success rate was far higher: of 96 designs produced in *E. coli*, 73 were expressed solubly (median soluble yield: 247 mg per liter of culture equivalent, Figure 3A) and 50 had the target monomeric or oligomeric state as assessed by SEC (Figure 3A,C). Many of the proteins were highly thermo-stable, with secondary structure being maintained up to 95 °C (Figure 3B).

**Fig. 3.**
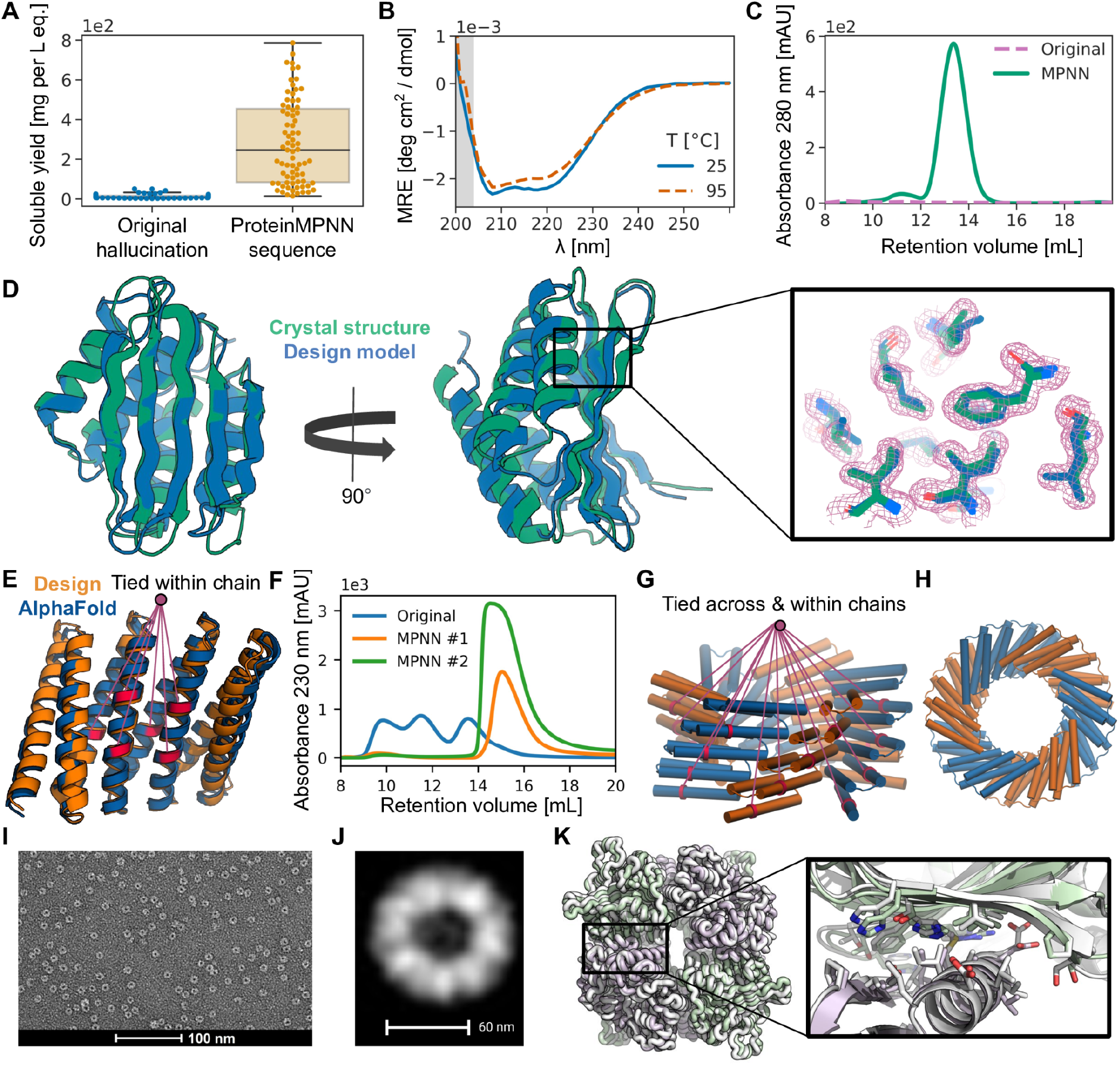
Structural characterization of ProteinMPNN designs. (**A**) Comparison of soluble protein expression over a set of AlphaFold hallucinated monomers and homo-oligomers (blue) and the same set of backbones with sequences designed using ProteinMPNN (orange), N=129. The total soluble protein yield following expression in *E. coli*, obtained from the integrated area unders size exclusion traces of nickel-NTA purified proteins, increases considerably from the barely soluble protein of the original sequences following ProteinMPNN rescue (median yields for 1 L of culture equivalent: 9 and 247 mg respectively). (**B**), (**C**), (**D**) In depth characterization of a monomer hallucination and corresponding ProteinMPNN rescue from the set in A. Like almost all of the designs in A, the sequence and structural similarity to the PDB of the design model are very low (E-value=2.8 against UniRef100 using HHblits, TM-score=0.56 against PDB). (**B**) The ProteinMPNN rescued design has high thermostability, with a virtually unchanged circular dichroism profile at 95 °C compared to 25 °C. (**C**) Size exclusion (SEC) profile of failed original design overlaid with the ProteinMPNN sequence design, which has a clear monodisperse peak at the expected retention volume. (**D**) Crystal structure of the ProteinMPNN (8CYK) design is nearly identical to the design model (2.35 RMSD over 130 residues), see Figure S5 for additional information. Right panel shows model sidechains in the electron density, in green crystal side chains, in blue AlphaFold side chains. (**E**), (**F**) ProteinMPNN rescue of Rosetta design made from a perfectly repeating structural and sequence unit (DHR82). Residues at corresponding positions in the repeat unit were tied during ProteinMPNN sequence inference. (**E**) Backbone design model and MPNN redesigned sequence AlphaFold model with tied residues indicated by lines (∼1.2Å error over 232 residues). (**F**) SEC profile of IMAC purified original Rosetta design and two ProteinMPNN redesigns. (**G**), **(H**) Tying residues during ProteinMPNN sequence inference both within and between chains to enforce both repeat protein and cyclic symmetries. (**G**) Side view of design model. A set of tied residues are shown in red. (**H**) Top-down view of design model. (**I**) Negative stain electron micrograph of purified design. (**J**) Class average of images from I closely match top down view in H. (**K**) Rescue of the failed two-component Rosetta tetrahedral nanoparticle design T33-27 (13) by ProteinMPNN interface design. Following ProteinMPNN rescue, the nanoparticle assembled readily with high yield, and the crystal structure (grey) is very nearly identical to the design model (green/purple) (backbone RMSD of 1.2 Å over two complete asymmetric units forming the ProteinMPNN rescued interface).

We were able to solve the X-ray crystal structure of one of the ProteinMPNN monomer designs with a fold more complex (TM-score=0.56 against PDB) than most *de novo* designed proteins (Figure 3D). The alpha-beta protein structure contains 5 beta strands and 4 alpha helices, and is close to the design target backbone (2.35 Å over 130 residues), demonstrating that ProteinMPNN can quite accurately encode monomer backbone geometry in amino acid sequences. The accuracy was particularly high in the central core of the structure, with sidechains predicted using AlphaFold from the ProteinMPNN sequence fitting nearly perfectly into the electron density (Figure 3D). Crystal structures and cryo-EM structures of ten cyclic homo-oligomers with 130-1800 amino acids were also very close to the design target backbones (accompanying manuscript, *Wicky et al*.). Thus, ProteinMPNN can robustly and accurately design sequences for both monomers and cyclic oligomers.

We next took advantage of the flexible decoding order of ProteinMPNN to design sequences for proteins containing internal repeats, tying the identities of proteins in equivalent positions. We found that many previously suboptimal Rosetta designs of repeat protein structures could be rescued by ProteinMPNN redesign, an example is shown in Figure 3E, F.

We next experimented with enforcing both the cyclic and internal repeat symmetry by tying positions both within and between subunits, as illustrated in Figure 3G. We experimentally characterized a set of C5/C6 cyclic oligomers built with Rosetta with sequences designed with Rosetta, and a second set with sequences designed using ProteinMPNN, and again observed much higher success rates with ProteinMPNN design. For the Rosetta designed set, 40% were soluble and none had the correct oligomeric state confirmed by SEC-MALS. For the ProteinMPNN designed set, 88% were soluble and 27.7% had the correct oligomeric state. We characterized the structure of one of the designs with sufficient size for resolution of structural features by negative stain EM (Figure 3I), and image averages were closely consistent with the design model (Figure 3J).

We next evaluated the ability of ProteinMPNN to design sequences that assemble into target protein nanoparticle assemblies. We started with a set of previously described protein backbones for two-component tetrahedral designs generated using a compute-and effort-intensive procedure that involved Rosetta sequence design followed by more than a week of manual intervention to eliminate unwanted substitutions (*15*). We used ProteinMPNN to design 76 sequences spanning 27 of these tetrahedral nanoparticle backbones, tying identities at equivalent positions in the 12 copies of each subunit in the assemblies, and ordered plasmids encoding them without further intervention. We found upon expression in *E. coli* and purification by SEC that 13 designs formed assemblies with the expected MW (∼1 MDa) (Figure S4). Although a similar overall success rate was obtained using Rosetta in the original study, several new tetrahedral assemblies were successfully generated using ProteinMPNN that had failed using Rosetta. We solved the crystal structure of one of these, and found that it was very close to the design model (1.2 Å Cɑ RMSD over two subunits, Figure 3K). Thus ProteinMPNN can robustly design sequences that assemble into designed nanoparticles, which have proven useful in several biotechnological applications including structure-based vaccine design (*16-18*). Sequence generation with MPNN is fully automated and requires only ∼1 second per backbone, vastly streamlining the design process compared to the earlier Rosetta-based procedure.

As a final test, we evaluated the ability of ProteinMPNN to rescue previously failed designs of new protein functions using Rosetta. We chose as a challenging example the design of proteins scaffolding polyproline helix motifs recognized by SH3 domains, such that portions of the protein scaffold outside of the SH3 peptide motif make additional interactions with the target, with the longer range goal of generating protein reagents with high affinity and specificity for individual SH3 family members. Backbones scaffolding a proline rich peptide recognized by the Grb2 SH3 domain were generated using Rosetta remodel (see Figure 4 legend), but sequences designed for these backbones did not fold to structures binding Grb2 when expressed in *E. coli* (Figure 4B, the design problem is challenging as very few native proteins have proline rich secondary structure elements that closely interact with the core of the protein). To test whether ProteinMPNN could overcome this problem, we generated sequences for the same backbones and expressed the proteins in *E. coli*. Biolayer interferometry experiments showed strong binding to the Grb2 SH3 domain (Figure 4B), with considerably higher signal than the free proline rich peptide; point mutations predicted to disrupt the design completely eliminated the binding signal. Thus ProteinMPNN can generate sequences for challenging protein design problems even when traditional RosettaDesign fails.

**Fig. 4.**
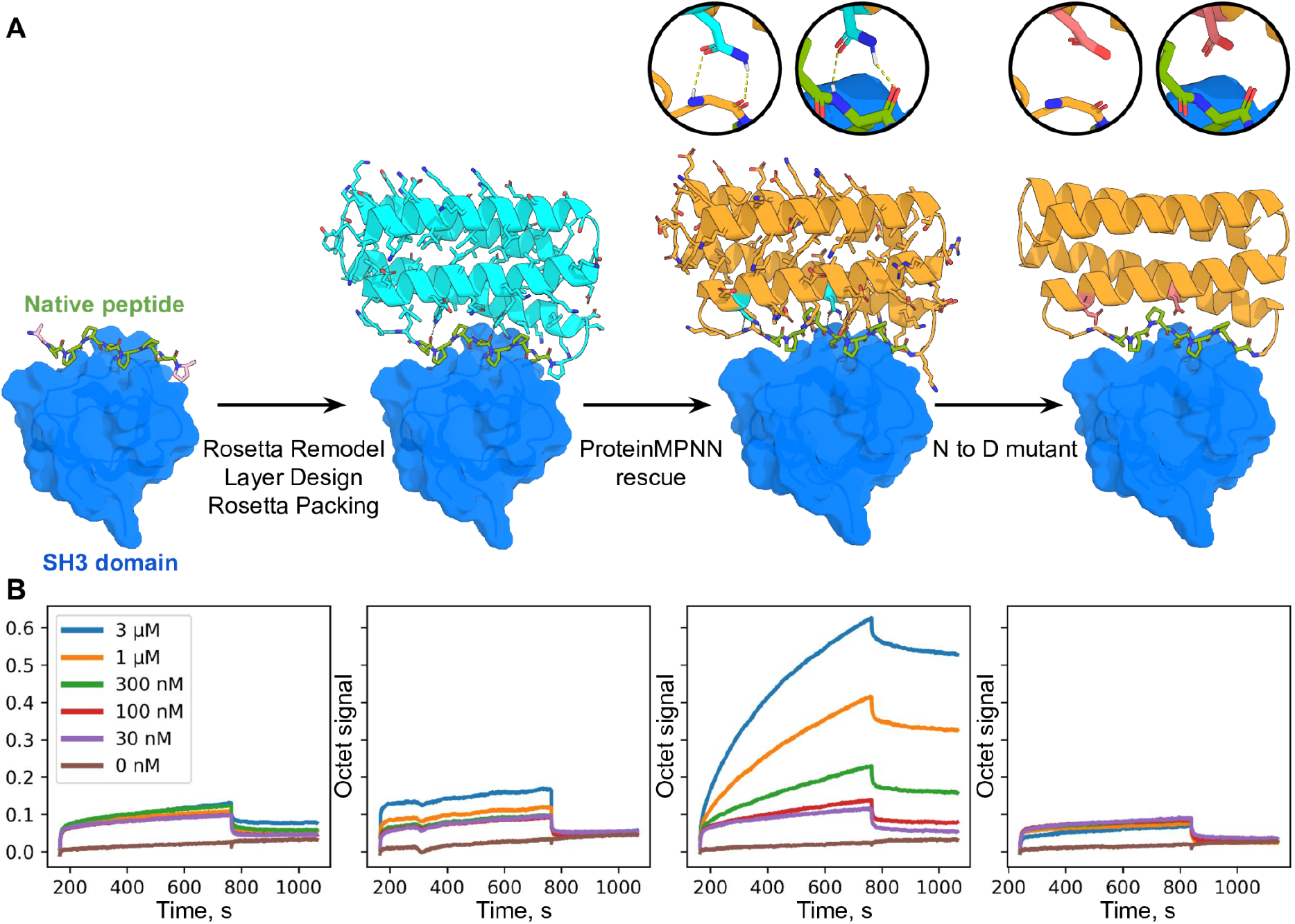
Design of protein function with ProteinMPNN. (**A**) Design scheme. First panel; structure (PDB 2W0Z) of the peptide APPPRPPKP bound to the human Grb2 C-term SH3 domain (peptide is in green, target in surface and colored blue). Second panel: helical bundle scaffolds were docked to the exposed face of the peptide using RIFDOCK (*19*), and Rosetta remodel was used to build loops connecting the peptide to the scaffolds. Rosetta sequence design with layer design task operations was used to optimize the sequence of the fusion (Cyan) for stability, rigidity of the peptide-helical bundle interface, and binding affinity for the Grb2 SH3 domain. Third panel; ProteinMPNN redesign (orange) of the designed binder sequence; hydrogen bonds involving asparagine sidechains between the peptide and base scaffold are shown in green and in the inset. Fourth panel; Mutation of the two asparagines to aspartates to disrupt the scaffolding of the target peptide. (**B**) Experimental characterization of binding using biolayer interferometry. Biotinylated C-term SH3 domain from human Grb2 was loaded onto Streptavidin (SA) Biosensors, which were then immersed in solutions containing varying concentrations of the target peptide (left) of the designs (right panels), and then transferred to buffer lacking added protein for dissociation measurements. The MPNN design (3rd panel from the left) has much greater binding signal than the original Rosetta design (2nd panel from the left); this is greatly reduced by the asparagine to aspartate mutations (last panel).

## Conclusion

ProteinMPNN solves sequence design problems in a small fraction of the time (1.2 sec vs 258.8 sec on a single CPU for a 100 residue protein) required for physically based approaches such as Rosetta, which carry out large scale sidechain packing calculations, achieves much higher protein sequence recovery on native backbones (52.4% vs 32.9%), and most importantly, rescues previously failed designs made using Rosetta or AlphaFold for protein monomers, assemblies, and protein-protein interfaces. Machine learning sequence design approaches have been developed previously (*1-6*), notably the previously described message passing method on which ProteinMPNN is based, but have focused on the monomer design problem, achieve lower native sequence recoveries, and with the exception of a TIM barrel design study (*6*) have not been extensively validated using crystallography and cryoEM to evaluate design accuracy. Whereas structure prediction methods can be evaluated purely in silico, this is not the case for protein design methods: In silico metrics such as native sequence recovery are very sensitive to crystallographic resolution (Figure S3 B, C) and may not correlate with proper folding (even a single residue substitution, while causing little change in overall sequence recovery, can block folding); in the same way that language translation accuracy must ultimately be evaluated by human users, the ultimate test of sequence design methods is experimental characterization.

Unlike Rosetta and other physically based methods, ProteinMPNN requires no expert customization for specific design challenges, and it should thus make protein design more broadly accessible. This robustness reflects fundamental differences in how the sequence design problem is framed. In traditional physically based approaches, sequence design maps to the problem of identifying an amino acid sequence whose lowest energy state is the desired structure. This is, however, computationally intractable as it requires computing energies over all possible structures, including unwanted oligomeric and aggregated states; instead Rosetta and other approaches as a proxy carry out a search for the lowest energy sequence for a given backbone structure, and structure prediction calculations are required in a second step to confirm that there are no other structures in which the sequence has still lower energy. Because of the lack of concordance between the actual design objective and what is being explicitly optimized, considerable customization can be required to generate sequences which actually fold; for example in Rosetta design calculations hydrophobic amino acids are often restricted on the protein surface as they can stabilize undesired multimeric states, and at the boundary region between the protein surface and core there can be considerable ambiguity about the extent to which such restrictions should be applied. While deep learning methods lack the physical transparency of methods like Rosetta, they are trained directly to find the most probable amino acid for a protein backbone given all the examples in the PDB, and hence such ambiguities do not arise, making sequence design more robust and less dependent on the judgement of a human expert.

The high rate of experimental design success of ProteinMPNN, together with the high compute efficiency, applicability to almost any protein sequence design problem, and lack of requirement for customization has made it the standard approach for protein sequence design at the Institute for Protein Design and we expect it to be rapidly adopted throughout the community. As illustrated in the accompanying paper (*Wicky et al*.), ProteinMPNN designs also have a much higher propensity to crystallize, greatly facilitating structure determination of designed proteins. The observation that ProteinMPNN generated sequences are predicted to fold to native protein backbones more confidently and accurately than the original native sequences (using single sequence information in both cases) suggests that ProteinMPNN may be widely useful in improving expression and stability of recombinantly expressed native proteins (residues required for function would clearly have to be kept fixed). We are currently extending ProteinMPNN to protein-nucleic acid design and protein-small molecule design which should increase its utility still further.

## Acknowledgements

The authors wish to thank Sergey Ovchinnikov, Chris Norn, David Juergens, Jue Wang, Frank DiMaio, Ryan Kibler, Minkyung Baek, Sanaa Mansoor, Luki Goldschmidt, and Lance Stewart for helpful discussions. The authors would also like to thank the Meta AI protein team for sharing AlphaFold models generated for UniRef50 sequences. The Berkeley Center for Structural Biology is supported in part by the National Institutes of Health (NIH), National Institute of General Medical Sciences. Crystallographic data collected at The Advanced Light Source (ALS) and is supported by the Director, Office of Science, Office of 20 Basic Energy Sciences and US Department of Energy under contract number DE-AC02-05CH11231.

## Author contributions

Conceptualization: JD, LFM, BIMW, AC, RJdH, HB, NBen

Methodology: JD, IA, PJYL

Software: JD, TFH, DT, BK, FC

Validation: JD, NBen, HB, AKB, BS, AK, HN, SP, PJYL, NBeth, RJdH, LFM, BIMW, AC, RJR

Formal analysis: JD, LFM, BIMW, RJR, NBen

Resources: JD, DB

Data curation: IA, JD, HB,

Writing – original draft: JD, DB

Writing – review & editing: JD, DB

Visualization: JD, RJR, RJdH, HB, LFM, BIMW, PJYL, HB

Supervision: DB, NPK

Project administration: JD

Funding acquisition: JD, DB

## Data and materials availability

All data is available in the main text or as supplementary materials. ProteinMPNN code is available at https://github.com/dauparas/ProteinMPNN.

## Supplementary Materials

### Materials and Methods

#### Methods for training single chain models

##### Training data

For single chain experiments presented in Table 1 we used a dataset based on the CATH 4.2 40% non-redundant set of proteins (*1, 7*). We trained models following the setup described in (*1*), i.e. using the learning rate schedule and initialization of the original Transformer paper (*20*), a dropout rate of 10% (*21*), a label smoothing rate of 10% (*22*), batch size with 6000 tokens, graph sparsity was set to be 30 nearest neighbors using Ca-Ca distances.

##### Architecture modifications

For experiment 1 we added extra input edge features, namely 16 Gaussian radial basis functions (RBFs) equally spaced from 0Å to 20Å for distances between N, Ca, C, O, and virtual Cb for i and j residues. This resulted in 25*16=400 edge features. The virtual Cb coordinates were calculated using ideal angle and bond length definitions: b = Ca -N, c = C - Ca, a = cross(b, c), Cb = -0.58273431*a + 0.56802827*b - 0.54067466*c + Ca.

For experiment 2 we introduced edge updates for the encoder network. The inputs to the encoder are node (denoted V_i_) and edge (denoted E_ij_) features for i and j residues. A message M_ij_ is constructed using a multilayer perceptron (MLP) applied to [V_i_, V_j_, E_ij_] concatenated tensors. These messages are summed over neighbors, j, and an additional MLP is applied to get a new updated nodes V_i_^new^. These new nodes are used to get new edges, E_ij_^new^=MLP[V_i_^new^, V_j_^new^, E_ij_]. We used layer normalization, dropout, and residual connections for all layers, h_new_ = LayerNorm[h_old_ + Dropout(dh)], where dh is an output from the layer, h_old_ is the old value, h_new_ is the updated new value.

For experiment 4 we implemented a random decoding order when training. To achieve this we constructed a random permutation matrix on-the-fly for every input example and applied it to rows and columns of the upper triangular matrix which is an autoregressive mask for the left to right decoding. Alternatively, one could permute input tokens and keep the autoregressive mask fixed given that the neural network architecture is permutation equivariant which is the case for most graph neural networks.

### Methods for training multi chain models

#### Training data

We trained ProteinMPNN on protein assemblies in the PDB (as of Aug 02, 2021) determined by X-ray crystallography or cryoEM to better than 3.5Å resolution and with less than 10,000 residues. Sequences were clustered at 30% sequence identity cutoff using mmseqs2 (*10*) resulting in 25,361 clusters. We split those clusters randomly into three groups for training (23,358), validation (1,464), and testing (1,539), ensuring that none of the chains from the target chain or chains from the biounits of the target chain would be in the other two groups. Every training epoch, we cycled through the sequence clusters and picked a random sequence member from each cluster, and for each such ‘query’ we randomly picked a protein conformation (in cases where there were multiple) and reconstructed the biological assembly for the corresponding PDB entry; for cases with multiple biological assemblies, we picked one assembly at random. For hetero-oligomeric assemblies, we masked out the sequence from the query chain but provided the network with the sequence information on all other chains in the assembly, while for homo-oligomers, sequences were masked out from all copies to prevent potential information leakage (two protein chains were considered as homo-oligomeric if the sequence identity between residues aligned by TM-align (*23*) was higher than 70%.

#### Loss function and optimization

We used negative log likelihood with a label smoothing rate of 10% (*22*) for the loss (not using label smoothing works well too). The sum of negative probabilities worked much better than the average of log probabilities. The training loss was defined by loss_average_ = sum(loss * mask) / 2000 where 2000 was chosen empirically, loss (categorical cross entropy per token) and mask had shapes [batch, protein length]. For optimization we used Adam with beta1 = 0.9, beta2 = 0.98, epsilon= 10−9, and the learning rate schedule described in (*20*). Models were trained using pytorch (*24*), batch size of 10k tokens, automatic mixed precision, and gradient checkpointing on a single NVIDIA A100 GPU. Training and validation losses (perplexities) as functions of optimizer steps are shown in Figure 3D. Validation loss converged after about 150k optimizer steps which is about 100 epochs of on-the-fly sampled training data from 23,358 PDB clusters.

#### Input features

ProteinMPNN input features were just embedded edges without any node features (Figure 1A). Using protein dihedral angles as node input features did not result in better performance so for simplicity in dealing with multichain backbones we did not use any node input features. The edge features consisted of distances between residues in Euclidean space and distances between residues in the primary sequence space (relative positional encoding) within a chain plus an indicator if residues are in different chains. We encoded distances between N, Ca, C, O, and virtual Cb (see Methods for training single chain models for the definition of Cb) for i and j residues using 16 RBFs equally spaced from 2Å to 22Å. For relative positional encoding we used AlphaFold (*9*) like discrete (one-hot encoded) tokens -32, -31,…, 31, 32 within the protein chains and additional token 33 if residues are in different chains. Ablating positional encodings showed almost the same performance suggesting that relative primary sequence or inter-chain information is already present in the Euclidean distances between atoms, e.g. distances between neighboring Ca atoms are the same for all residues.

#### Model architecture

We used encoder-decoder message passing neural networks for this task (*1, 20, 25*), see Figure 1. The encoder takes graph nodes and edges as inputs and using 3 layers with hidden dimension of 128 (larger hidden dimensions mainly decrease training loss with only marginal gains to the validation loss) updates those nodes and edges using message passing with edge updates.

Pseudocode for the encoder layer (V - node features, E - edge features):

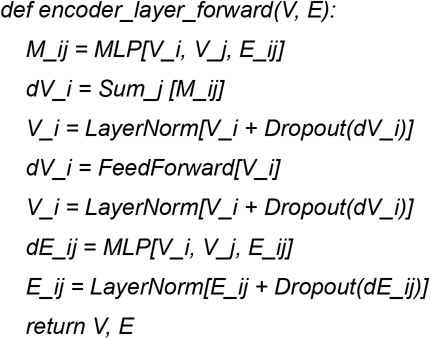

The decoder takes in node and edge features from the encoder plus encoded protein sequence to which an autoregressive mask is applied. The decoder layer is a vanilla MPNN layer with 3 layers and 128 hidden dimensions.

Pseudocode for the decoder layer (V - node features, E - edge features, S - sequence features, mask - autoregressive mask):

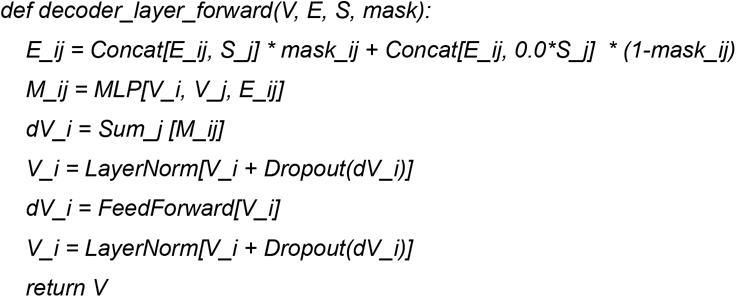

For both encoder and decoder the multi-layer perceptrons (MLPs) had 3 linear layers of the model’s hidden dimension and GELU activation functions (GELU worked slightly better compared with ReLU). The FeedForward layers had 2 linear layers with GELU activations and with the middle hidden dimension 4 times bigger than the input dimension as in the Transformer paper (*20*).

### Further in silico analysis

#### Amino acid compositional bias

Figure S2 shows ProteinMPNN and Rosetta amino acid compositional bias compared with the native PDB sequences. ProteinMPNN sequences were designed using temperature T=0.1. Rosetta designed sequences have an overrepresentation of alanines in the core and boundary. Both models favor a negatively charged glutamic acid, E, on the surface and disfavor a polar amino acid glutamine, Q. Interestingly, ProteinMPNN and Rosetta biases strongly disagree for lysine, K on the surface. Amino acid bias for ProteinMPNN is a function of the sampling temperature (see Figure S6) with low temperatures introducing more charged amino acids on the surface.

#### AlphaFold benchmarks

We ran all 5 AlphaFold ptm models with 3 recycles and selected the model with the highest average pLDDT as described in the AlphaFold paper (*9*) using only a single sequence as an input. For results described in Figure 2C and Figures S7, S8, S9, we generated 8 sequences per target backbone for 396 monomers (with maximum length of 300 residues) from the test set and used AlphaFold to predict structures for these sequences. Dependence on the inference noise level and sampling temperature for sequence recovery and relative AlphaFold success rate for the MPNN model trained with 0.2Å are shown in Figure S7A, B. Both of these metrics decrease with higher levels of noise and temperature. We also generated 8 sequences per target backbone for 277 homomers (with maximum length of 400 residues) and plotted sequence recovery and inter-chain predicted aligned error (PAE) as a function of the MPNN training noise level, Figure S7C. The trend is very much the same as for the monomer case showing that sequences from slightly noised MPNN models are more easily decoded by AlphaFold. It is important to notice that AlphaFold success rate for the single sequence prediction depends on the number of recycles used during the inference. We benchmarked 1, 2, 3, 6, and 12 recycles for the MPNN monomer sequences, see Figure S7D. The success rate monotonically increases suggesting that more powerful single sequence structure prediction models might have even higher success rates, i.e. MPNN sequences are correctly encoding structures, but it is hard to predict those structures using only single sequence information. Finally, we looked at the dependence between AlphaFold pLDDT and true lDDT-Ca, i.e. between native target backbone for MPNN and predicted AlphaFold backbone, Figure S8. The correlation is very much like in the original AlphaFold paper (*9*) in this single sequence regime with slight underestimation of lDDT-Ca.

#### Ca-only ProteinMPNN

We trained ProteinMPNN which used Ca coordinates only as an input instead of full backbone coordinates to have a way to generate sequences for coarse, or approximately correct backbones. In the similar way as for the full atom ProteinMPNN adding inter-atom distances as edge features helped to improve model’s performance. We added distances between triplets of Ca atoms: Ca_i-1_, Ca_i_, Ca_i_ and Ca_j-1_, Ca_j_, Ca_j_ for residues i and j encoded as 16 Gaussian radial basis functions. The rest of the architecture is the same as for the full atom version. Sequence recovery and relative AlphaFold success rate as a function of training noise level are shown in Figure S9.

### Experimental methods

LM0878 (Figure 1D) was expressed in TBM-5052 medium with 50 μg/mL Kanamycin for 24 hours at 37°C with shaking at 225 rpm using the autoinduction method before harvesting via centrifugation at 4000xg for 5 minutes. Cells were lysed in 30ml of wash buffer (20 mM Tris, 300 mM NaCl, 25 mM Imidazole, pH 8.0) with sonication at 4°C. Lysates were then centrifuged for 45 minutes at 14000xg and applied to Ni-NTA resin that was pre-equilibrated with wash buffer. The resin was washed with 30 column volumes of wash buffer. 6xhis affinity tags were removed via on-bead SNAC (*26*) cleavage. Resin was washed with 20CV of cleavage buffer (100 mM CHES, 100 mM Acetone oxime, 100 mM NaCl, pH 8.6) prior to incubation with 20ml cleavage buffer and 2mM NiCl2 overnight at room temperature. Flow through was collected and concentrated to 1ml using 3K protein concentrators (Millipore Sigma) before size exclusion chromatography using an S75 10/300 GL increase column (GE Healthcare).

LM0878 was crystallized through sitting drop vapour diffusion at room temperature in 0.1M citric acid pH 3.5 and 3M sodium chloride. Prior to harvesting and flash freezing in liquid nitrogen, crystals were transferred to the crystallization condition with 25% ethylene glycol. Diffraction data was collected at 100K at the Advanced Light Source beamline 8.2.1. Images were integrated using XDS 20220110 (*27*), with Aimless (*28*) used for scaling and merging. The design model was used as the search model for molecular replacement with Phaser 2.8. *(29*) Model building and refinement was done using Coot 0.9.8 (*30*), and Phenix refine from Phenix 1.20. (*31*) All structures were validated using MolProbity 4.5.1. (*32*) Crystallographic statistics are available in Table S1. Crystallographic data has been deposited to the protein bank with the PDB ID 8CYK.

**Fig. S1.**
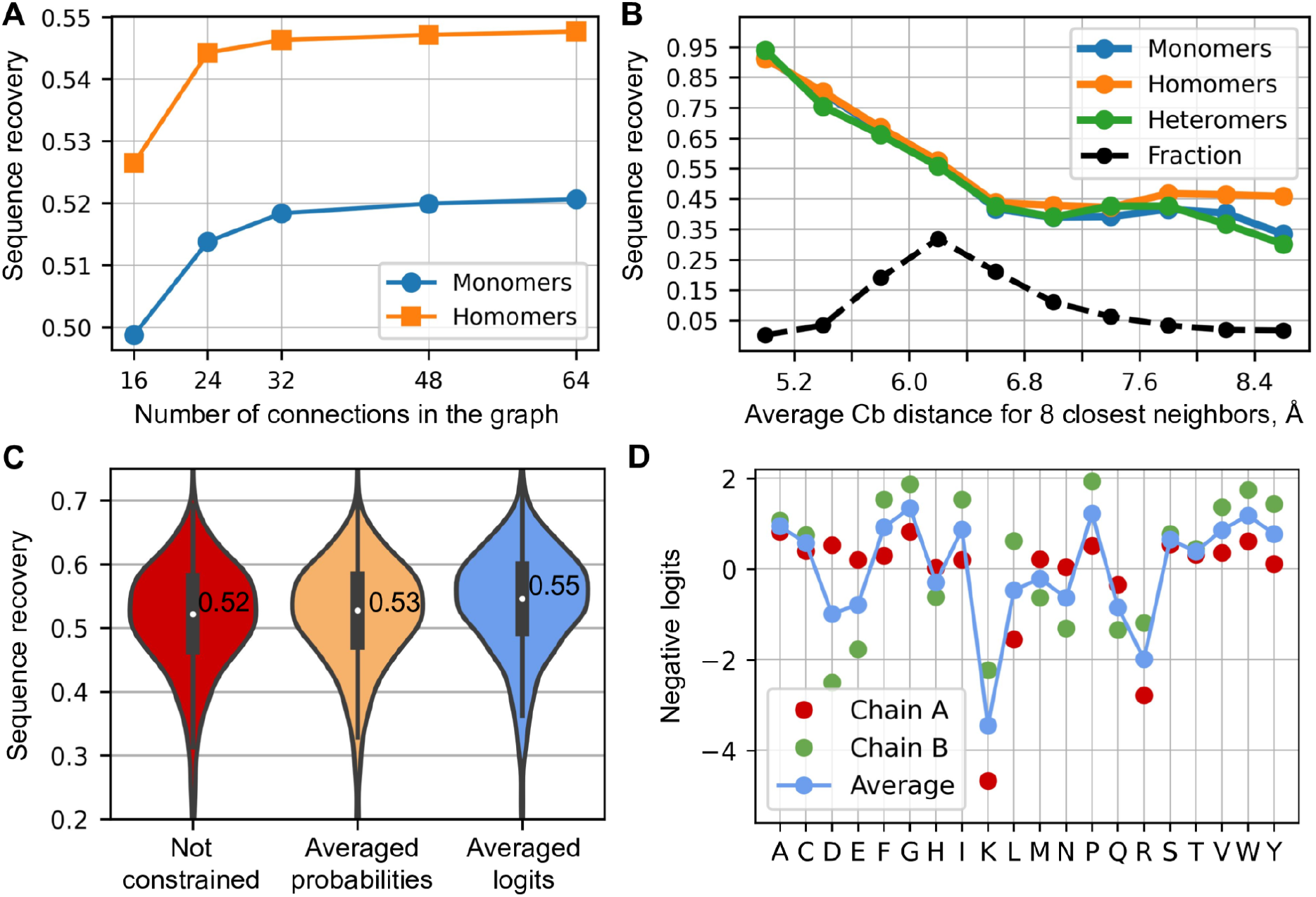
In silico validation results. (**A**) Sequence recovery as a function of the number of nearest neighbors in the graph. (**B**) Sequence recovery as a function of burial for monomers, homomers, and heteromers. (**C**) Comparing three different ways of generating sequences for homomers: unconstrained (treating as non-symmetric), averaging predicted probabilities, averaging predicted logits. (**D**) An example showing negative logits predicted by the model for chain A and chain B in the homodimer. The blue curve shows averaged logits which will be normalized to sample an amino acid.

**Fig. S2.**
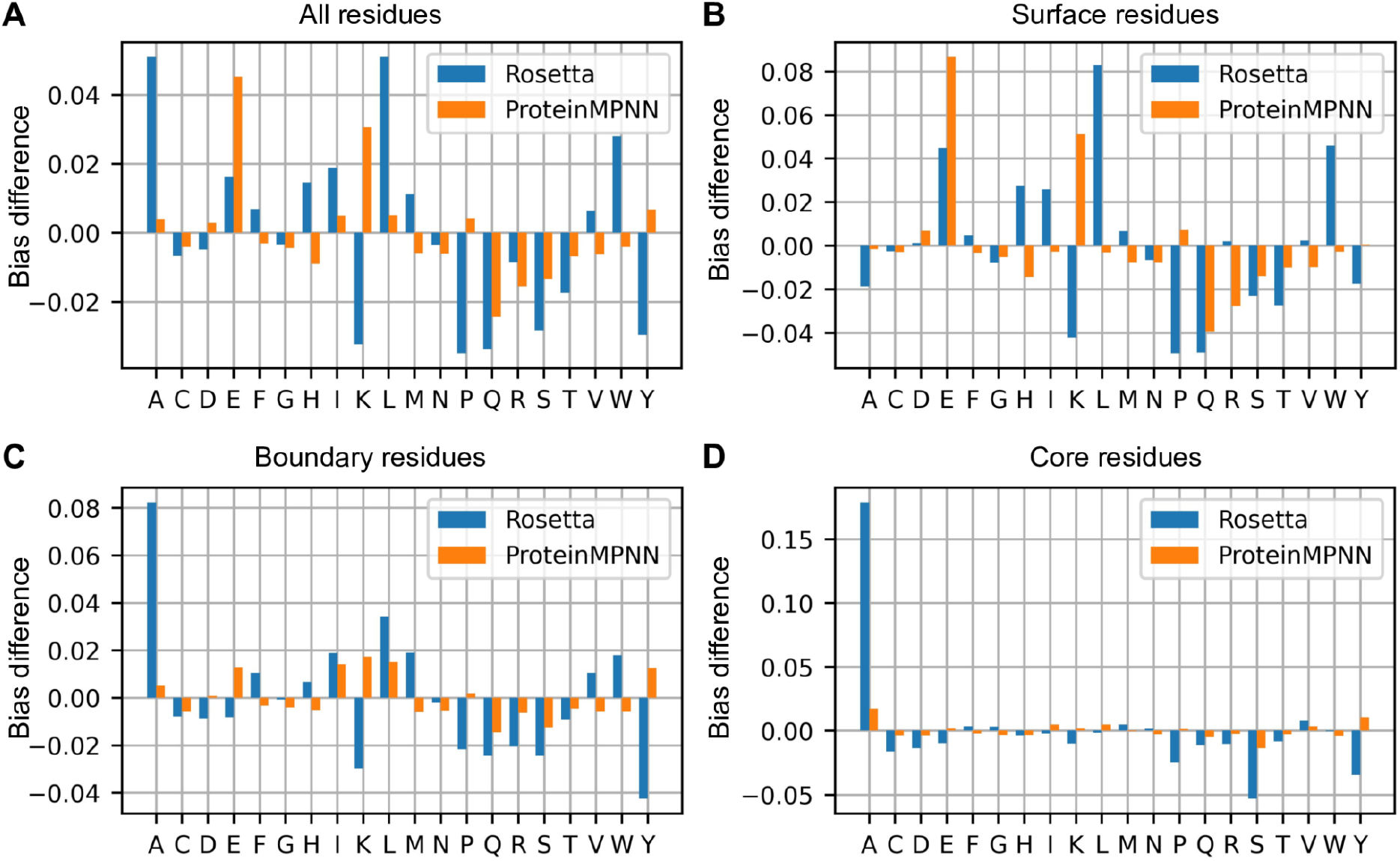
Difference in compositional bias for Rosetta and ProteinMPNN. (**A**) Bias for all residues in the monomer chain. (**B**) Bias for surface residues. (**C**) Bias for boundary residues. (**D**) Bias for core residues.

**Fig. S3.**
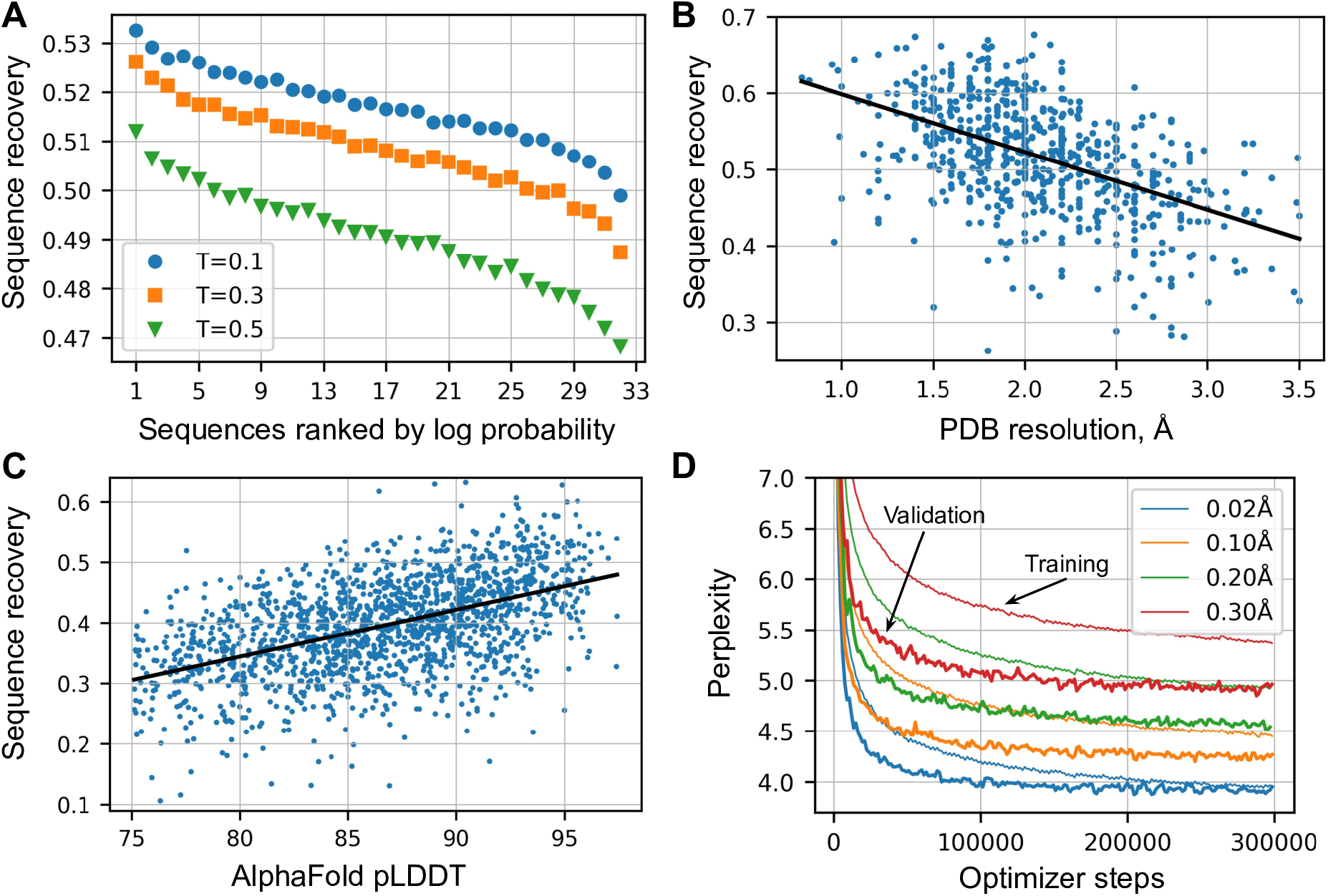
Sequence recovery dependence on confidence and backbone quality. (**A**) We generated 32 sequences for every backbone in the test set of 690 PDB monomers using sampling temperatures 0.1, 0.3, 0.5 and ranked them using the log probability of the model (confidence). (**B**) Sequence recovery as a function of PDB resolution using 0.02Å noised MPNN model for a set of 690 PDB monomers. The black line shows a least squares linear fit. Spearman’s rank correlation was -0.487. (**C**) Sequence recovery as a function of AlphaFold model confidence (average pLDDT) for a set of 1621 UniRef50 models. Spearman’s rank correlation was 0.502. (**D**) Training and validation perplexities as a function of optimizer steps for different levels of backbone noise. Backbone noise and dropout (0.1) was not applied during the validation.

**Fig. S4.**
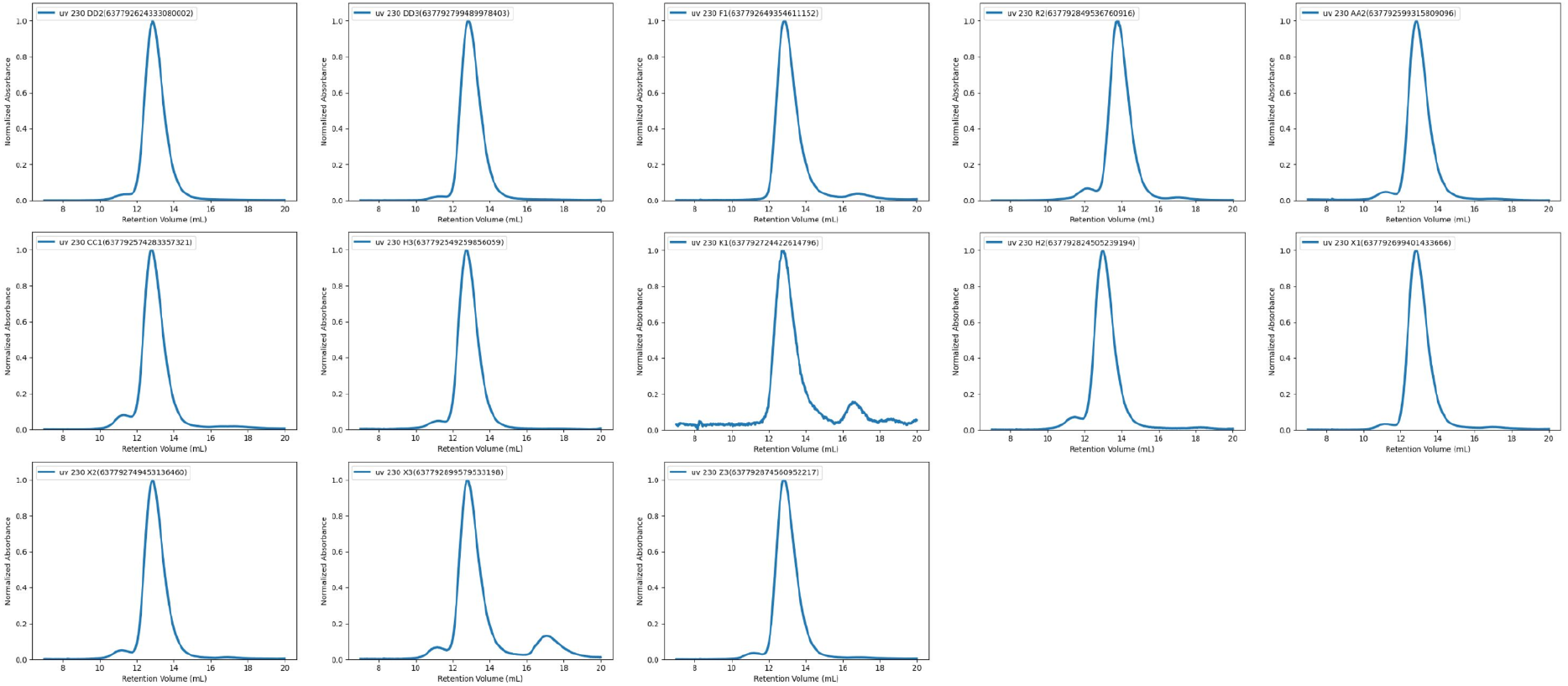
Size exclusion chromatography of proteinMPNN designed two-component tetrahedron nanoparticles. 13/76 nanoparticle designs eluted at ∼13 mL on a Superdex S200 10/300 Increase column, corresponding to nanoparticles of ∼1 MDa. On the y-axis is normalized absorbance (uv 230), on the x-axis is retention volume [mL].

**Fig. S5.**
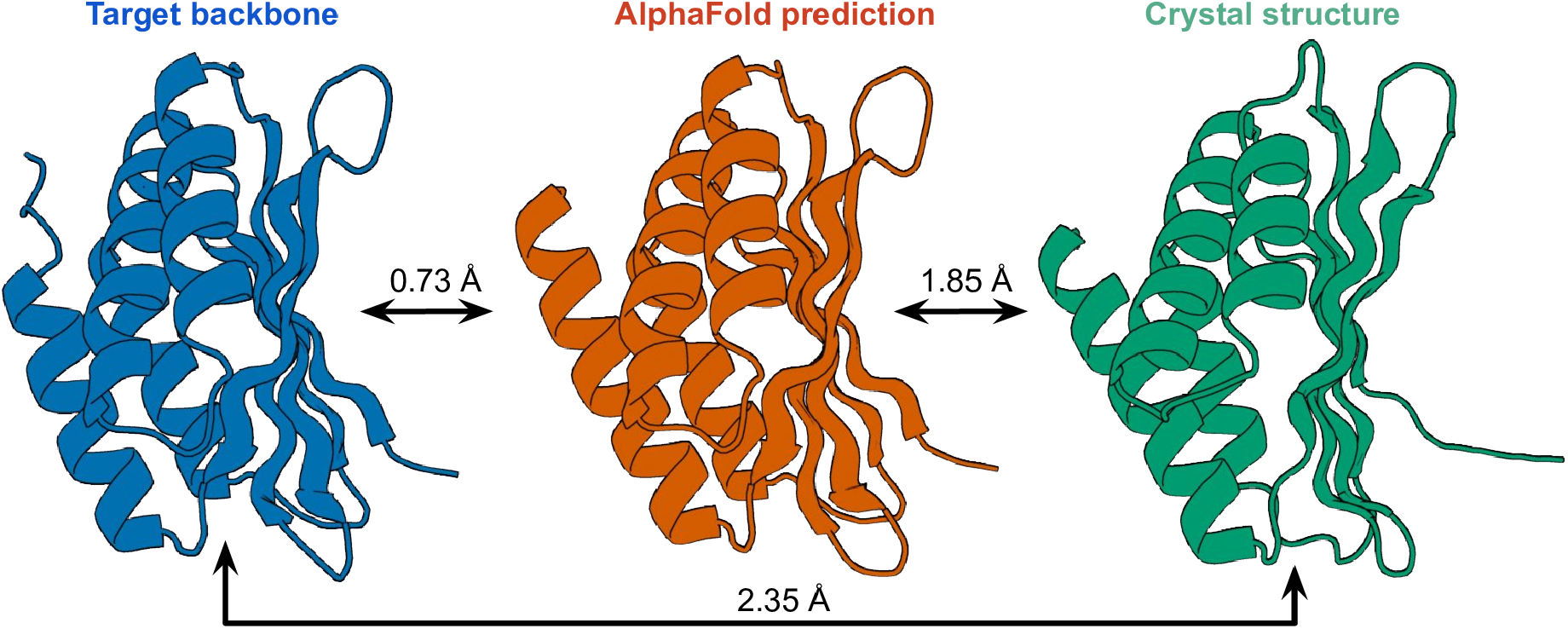
Designed backbone and crystal structure comparison. Target backbone (LM0878) from hallucination on the left, AlphaFold prediction using ProteinMPNN sequence in the middle, and crystal structure on the right (deposited to PDB as 8CYK). Backbone RMSDs are shown for every pair of backbones.

**Fig. S6.**
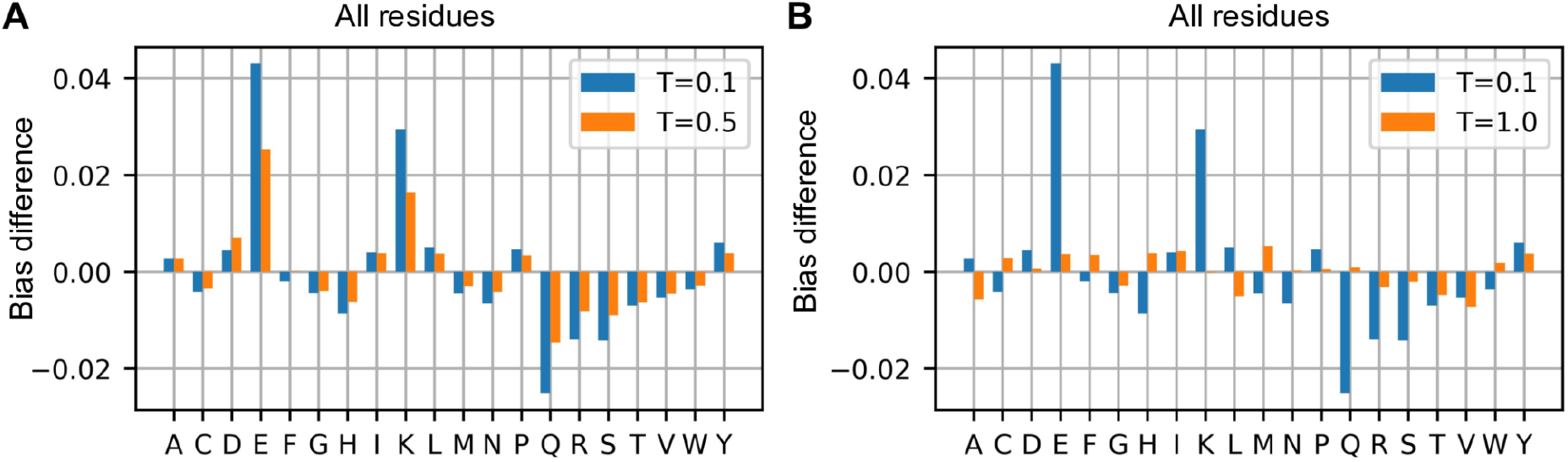
ProteinMPNN bias for different sampling temperatures. (**A**) ProteinMPNN generates more charged amino acids in expense of the polar ones at low temperatures which likely leads to highly thermo-stable proteins. (**B**) Amino acid bias is very small at temperature T=1.0.

**Fig. S7.**
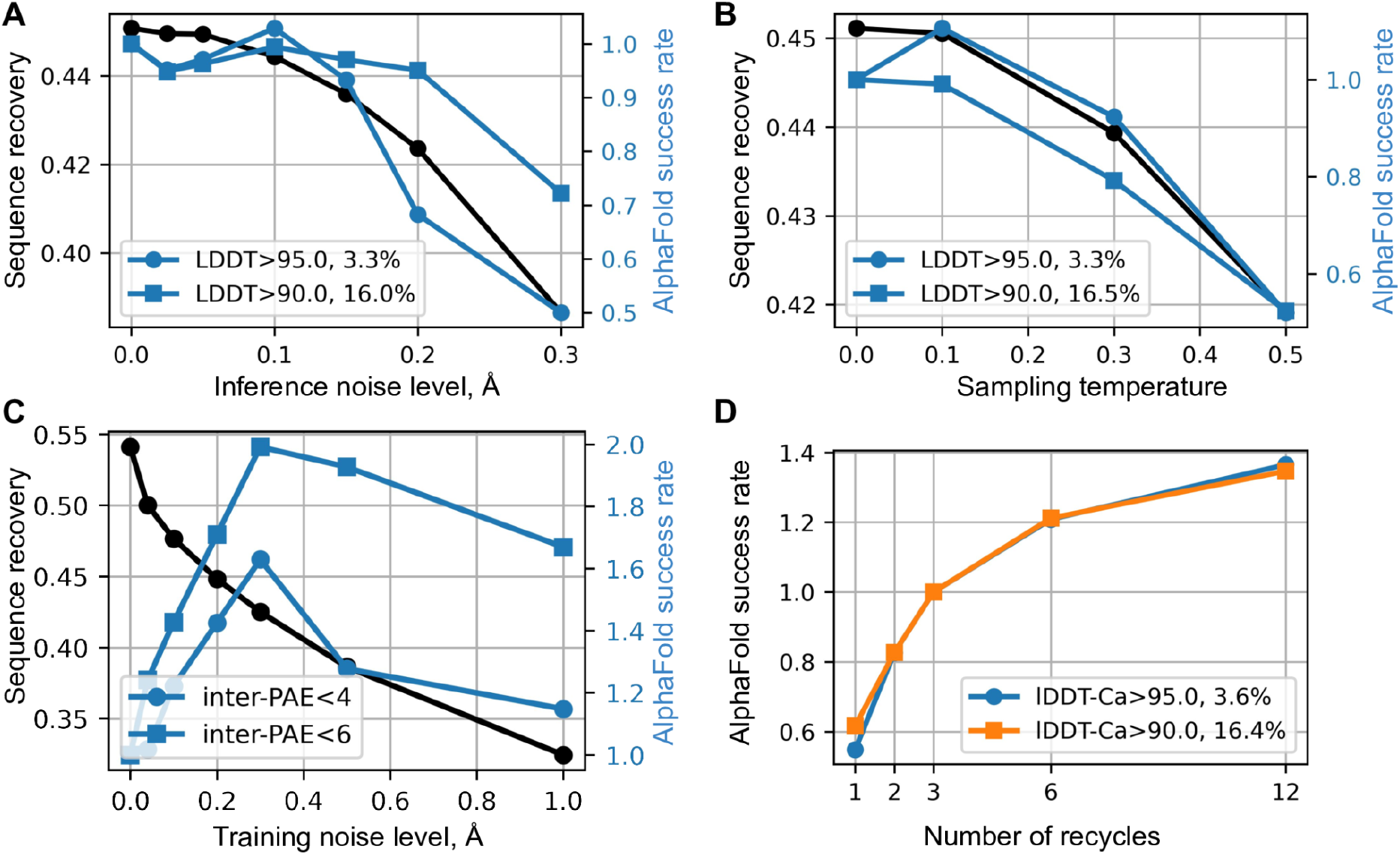
Sequence recovery and AlphaFold (AF) benchmarks for the ProteinMPNN trained with 0.2Å noise level. (**A**) Sequence recovery and AF success rate for monomeric structures with true LDDT>95.0, 90.0 as a function of noise applied to backbones during the inference, 1.0 corresponds to 3.3% absolute rate for 95.0 LDDT cutoff and to 16.0% for 90.0 cutoff. (**B**) Same as A, but as a function of sequence sampling temperature. (**C**) Sequence recovery and AF success rate for homomers with inter-chain PAE<4.0, 6.0 as a function of training noise level, 1.0 corresponds to 2.4% for the 4.0 PAE cutoff, and to 5.6% for the 6.0 cutoff. (**D**) AlphaFold success rate as a function of number of AlphaFold recycles for monomers.

**Fig. S8.**
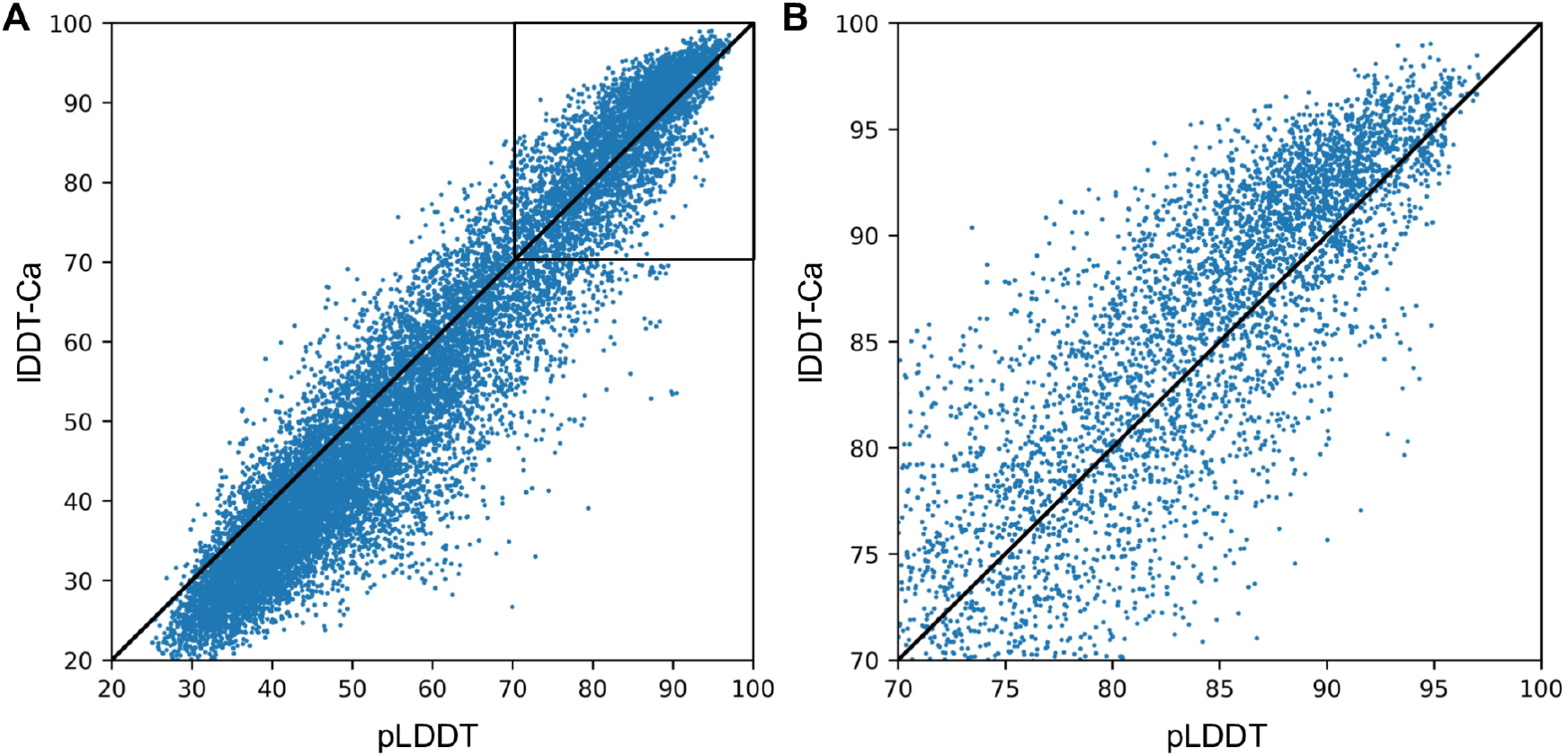
Comparing true lDDT-Ca with AlphaFold predicted pLDDT for MPNN sequence redesigns on native PDB backbone monomers using single sequence prediction. lDDT-Ca is calculated between the native backbone input to the MPNN and the AlphaFold output using MPNN sequence. (**A**) pLDDT and lDDT-Ca are highly correlated in this regime. (**B**) Zoomed in version of A showing that true lDDT-Ca is slightly underestimated by pLDDT.

**Fig. S9.**
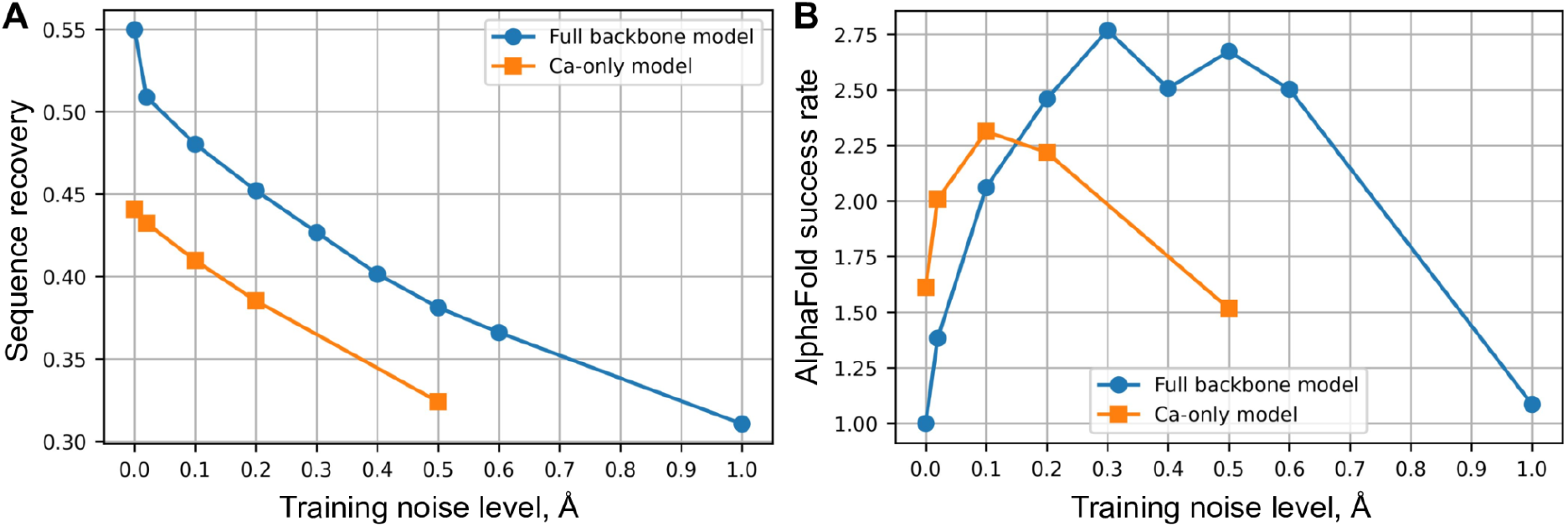
Benchmarks of Ca-only ProteinMPNN model for monomers. (**A**) Sequence recovery as a function of training noise level for Ca-only and full backbone models. (**B**) Relative AlphaFold success rate as a function of training noise level for Ca-only and full backbone networks for the models with lDDT-Ca>90.0, 1.0 is equal to 6.7% success rate.

**Table S1.** Crystallographic statistics for HALC1_878.

|  |  |
| --- | --- |
|  | **HALC1_878 (PDB ID: 8CYK)** |
| **Data Collection** |  |
| Space group | I 1 2 1 |
| Cell dimensions |  |
| a, b, c (Å) | 64.17, 55.45, 82.81 |
| $\alpha$, $\beta$, $\gamma$ (°) | 90.00, 94.00, 90.00 |
| Resolution (Å) | 48.98 - 1.65 (1.709 - 1.65) |
| Rmerge (%) | 2.1 (19.4) |
| Rpim (%) | 2.1 (19.4) |
| $\sigma(I)$ | 15.91 (3.96) |
| CC 1/2 | 0.999 (0.942) |
| Completeness (%) | 96.85 (95.70) |
| Redundancy | 2.0 (2.0) |
| **Refinement** |  |
| No. unique reflections | 33946 (3322) |
| Rwork / Rfree (%) | 18.5 (24.0) / 21.7 (28.1) |
| No. non-hydrogen atoms | 2366 |
| Macromolecules | 2151 |
| Water | 215 |
| Ramachandran favoured/allowed | 98.83/1.17 |
| **R.m.s. deviations** |  |
| Bond lengths (Å) | 0.011 |
| Bond angles (°) | 1.12 |
| **B-factors (Å<sup>2</sup>)** |  |
| Macromolecules | 28.36 |
| Solvent | 39.25 |

